# PyFuRNAce: An integrated design engine for RNA origami

**DOI:** 10.1101/2025.04.17.647389

**Authors:** Luca Monari, Ina Braun, Erik Poppleton, Kerstin Göpfrich

## Abstract

To realize the full potential of RNA nanotechnology and RNA origami, user-friendly design tools are needed. Here, we present pyFuRNAce, an open-source, Python-based software package with a graphical user interface that enables the design of complex RNA nanostructures. PyFuRNAce integrates the entire RNA origami workflow—from motif definition and blueprint design to sequence generation and primer selection—into a single, user-friendly platform. Built around a motif-based assembly paradigm, the software enables users to create and modify custom nanostructures through an intuitive web interface with streamlined design steps and real-time 3D visualization. By consolidating multiple design stages into a unified environment, pyFuRNAce reduces the entry barrier for RNA nanotechnology and accelerates the development of functional RNA origami structures for applications in medicine, biotechnology, and synthetic biology.

## Introduction

RNA is widely known as an intermediary between two seemingly more interesting classes of molecules: DNA, the information storage, and proteins, the functional building blocks of a cell. However, the last fifty years of molecular biology have been characterized by an increasing appreciation for the diverse roles that RNA plays in cellular function. Central to its diversity is the ability of RNA to fold into complex secondary, tertiary and quaternary structures—like proteins. Scientists, naturally, asked whether we would be able to design RNA sequences with defined structures and functions. The field which emerged from this question, RNA nanotechnology, is still in its infancy compared to protein design. This is not only due to the historical lack of interest, but also the hard-to-predict dynamic and flexible nature of RNA, and the fact that RNA is built from just four similar nucleotides, which interact in relatively simple and less predictable ways—unlike proteins, whose diverse amino acids form a wide range of interactions like hydrophobic cores and salt bridges to guide folding [1]. The number of experimentally-characterized RNA structures is a fraction of what is available for proteins (200,000 protein structures vs less than 2,000 RNA structures), making machine learning approaches difficult [1]. Yet recent years have revealed promising and wide-ranging applications of RNA nanotechnology in biotechnology, synthetic biology, material science and medicine, where aptamer-based drugs are reaching the clinical trial phase[2, 3]. Early advancements in RNA nanotechnology demonstrated the structural capabilities of RNA through rational design of discrete and periodic 2D and 3D shapes, such as sheets [4], nanotubes [5], 2D crystals [6], 3D crystals [7], and polyhedra [8]. To accomplish these feats of rational design, the field has developed three RNA design pardigms: RNA architectonics (tectoRNA) [9, 10], co-transcriptional RNA origami [4, 11], and paranemic cohesion (PX) RNA origami [12], each with their advantages and disadvantages. In addition to structures demonstrating our ability to control RNA, RNA nanostructures have also been used in numerous preliminary studies in medicine[13, 14], with applications including siRNA delivery [15], immunomodulation [16, 17], and protein inhibition [18, 19, 3].

Despite progress in RNA nanotechnology, it remains underdeveloped compared to the more established field of DNA nanotechnology [20, 21]. DNA nanotechnology has seen tremendous advancements, in particular, thanks to the DNA origami design paradigm [22], which enabled the production of large, arbitrary structures at high yield. Although the names are similar, co-transcriptional RNA origami has some unique features which may be advantageous over DNA origami for applications in medicine, biotechnology and synthetic biology. Co-transcriptional RNA origami can be produced in a single biochemical step without the need for thermal annealing, offering the potential for mass production. Additionally, RNA origami benefits from a large library of aptamers and ribozymes that can be used to directly integrate functional motifs without requiring covalent chemical modification, an advantage for direct production in cellular environments [2]. Considering the benefits of co-transcriptional RNA origami over DNA origami, the different maturities of the two fields are striking.

The limited development of RNA origami can largely be attributed to one key factor: the lack of user-friendly design tools. In contrast, DNA origami has seen the emergence of a variety of design tools, such as CaDNAno [23], Tiamat [24], ENSnano [25], Athena [26], SARSE [27] and MagicDNA [28], which have been crucial in the field’s rapid advancement. These tools have enabled diverse applications and even led to the establishment of companies focused on DNA origami, including its use in vaccine trials [29, 30].

As far as RNA design software is concerned, the landscape is relatively sparse. Tiamat was used to design PX RNA origami [12]. However, it would benefit from an easy way of drawing large structures and means to integrate non-helical structures. RNA architectonics has benefited from tools such as Assemble2 [31] which facilitate fragment-based modeling; however, most tools are primarily aimed at structure prediction and refinement, rather than nanostructure design.

Co-transcriptional RNA origami on the other hand, has a tailored design tool: the RNA Origami Automated Design (ROAD) software package [11]. ROAD is a command-line tool that employs Perl scripts to generate RNA origami structures from blueprint files (Trace analysis and Trace pattern), build 3D structures from a fixed motif library (RNAbuild), and perform sequence optimization through inverse folding and sequence symmetry minimization (Revolvr). However, ROAD relies on text-based schematic files that require precise character alignment through manual editing, which is time-consuming and error-prone. Furthermore, adding new motifs to ROAD’s motif library is non-trivial, making it difficult to take full advantage of the diversity of functional RNA motifs.

To streamline RNA origami design, we introduce pyFuRNAce. PyFuRNAce is a userfriendly, open-source web application. It provides an intuitive graphical user interface (GUI), real-time 3D visualization, and a complete interface for the entire RNA origami design pipeline. Moreover, pyFuRNAce integrates a flexible motif library, enabling users to select or create custom motifs to meet specific design needs. These features make RNA origami design accessible to experts and non-experts alike, empowering researchers to explore the vast potential of RNA origami.

## Materials and methods

PyFuRNAce was developed and tested using Python 3.12 (minimum supported version: 3.10). The dependencies include Streamlit [32] (v1.43) to build the user interface; and NumPy [33] (v2.2.2) and SciPy [34] (v1.14.1) to handle core functionality for 3D structures. oxDNA Analysis Tools [35] is an optional dependency that supports PDB file reading/writing and preparing designs for oxRNA simulations [36, 37].

Several external Streamlit components were used to build an interactive and responsive user interface, including st-click-detector [38] for interactive RNA origami blueprint editing, st oxview [39] for 3D structure visualization using oxView, and st forna component [40] for rendering secondary structures in dot-bracket notation via Forna [41] on the Generate page. PyFuRNAce includes and integrates Revolvr from the ROAD package [11] for RNA sequence generation. To support different operating systems, we modified the Revolvr script to rely on the ViennaRNA python API [42, 43], already required for pyFuRNAce, and not on a locally compiled copy of ViennaRNA.

In the end, the Biopython library [44] is used to implement sequence conversion, melting temperature calculations and primer design. While pyFuRNAce does not include the oxDNA simulation engine [45, 2], it generates all required input files for downstream molecular dynamics simulations using the oxRNA force field [36].

The pyFuRNAce package is cross-platform and compatible with Windows, macOS, and Linux operating systems. pyFuRNAce can be accessed at http://pyfurnace.de.

### RNA 3D structure generation

All 3D structures in pyFuRNAce are internally represented in the oxDNA format [46], (‘new topology’ format with 5’ to 3’ directionality). The 3D geometries of common structural elements—such as stems, dovetails, and 180° kissing loops, including their branched and dimeric forms—are computationally generated using OxView parameters [35] for ideal RNA A-form helices. Dovetail crossover geometries are adapted from the average coordinates of crossover points identified in the PDB structure 7QDU [47].

All other 3D motifs used in pyFuRNAce are either extracted from experimentally determined structures available in the Protein Data Bank (PDB IDs are listed in the code documentation) or predicted using RNA Composer [48, 49] when no experimental structure is available.

## Results

The primary goal of pyFuRNAce is to streamline the RNA origami design workflow – from motif definition, blueprint design and 3D visualization to sequence generation, template preparation and primer selection – within a unified and user-friendly interface. RNA origami design is often iterative; multiple refinement cycles are necessary to optimize the geometry, sequence and folding. PyFuRNAce supports this process by integrating all major design stages into a cohesive environment, replacing the previously fragmented pipeline that relied on multiple tools (e.g. text editors for blueprint creation, RNAbuild and ChimeraX for visualization, Revolvr and RNAfold for sequence generation and folding validation, separate utilities for DNA conversion, and external calculators for primer melting temperature). Additionally, pyFuRNAce integrates a robust Python library that allows users to design RNA origami blueprints through Python scripts or Jupyter notebooks. Users can automate the design process, generate RNA origami sequences, and customize the workflow. This capability enables the creation of complex RNA structures through code, facilitating large-scale design iterations, systematic optimization, and integration with other computational tools, offering advanced users complete control over the design process.

The user interface is organized into four modules: Design, Generate, Convert, and Prepare (Figure 1a), each corresponding to a distinct stage in the RNA origami design pipeline. In addition to merging the workflow into a single web application, pyFuRNAce introduces several key features that simplify and enhance RNA origami design (Figure 1b), including:

**Figure 1.**
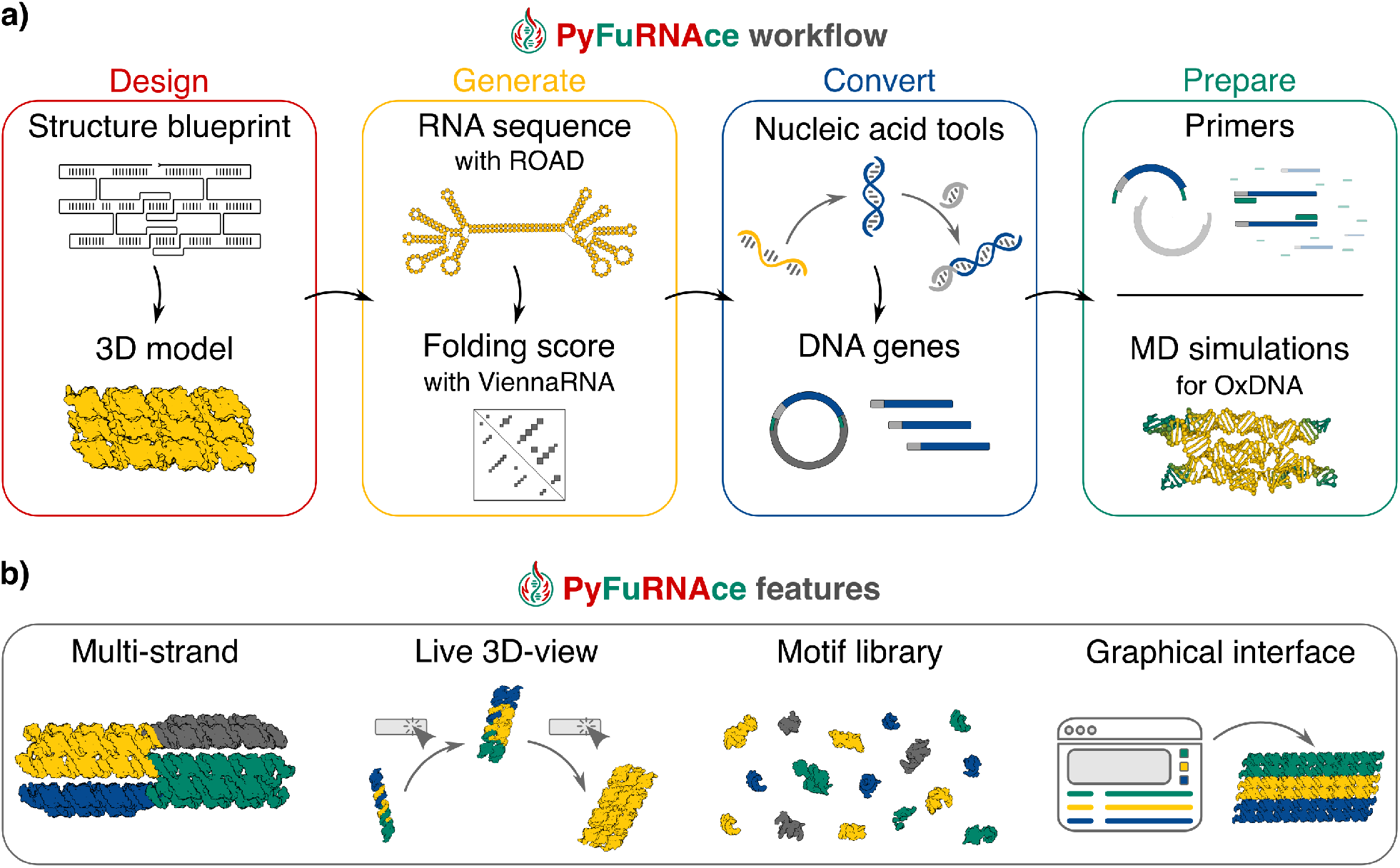
**a)** The RNA origami computational workflow integrated with pyFuRNAce. The Design page allows for blueprint and 3D structure design; the Generate page produces RNA sequences and structure folding scores; the Convert page analyzes the nucleic acid sequences and outputs a DNA template; the Prepare page helps in the primer design and the preparation of files for oxRNA MD simulations. PyFuRNAce can we accessed at http://pyfurnace.de. **b)** New features to streamline RNA origami design introduced with pyFuRNAce.

- Support for multi-strand design in the blueprint generation module, enabling users to visualize partial designs or multimeric assemblies.
- Real-time 3D visualization of RNA nanostructures.
- A rich and expandable motif library to keep the tool updated with the latest aptamer discoveries.
- A user-friendly graphical interface (GUI) that lowers the entry barrier to RNA origami design, with the option to extend functionality through an integrated Python scripting interface.

### Design

The Design module is the platform’s central feature. It enables object-oriented motif definition and blueprint generation through a graphical user interface (Figure 2). At the top of the interface, users can access and insert motifs from the pyFuRNAce library. Below, the RNA origami blueprint and the 3D structure are displayed with interactive components.

**Figure 2.**
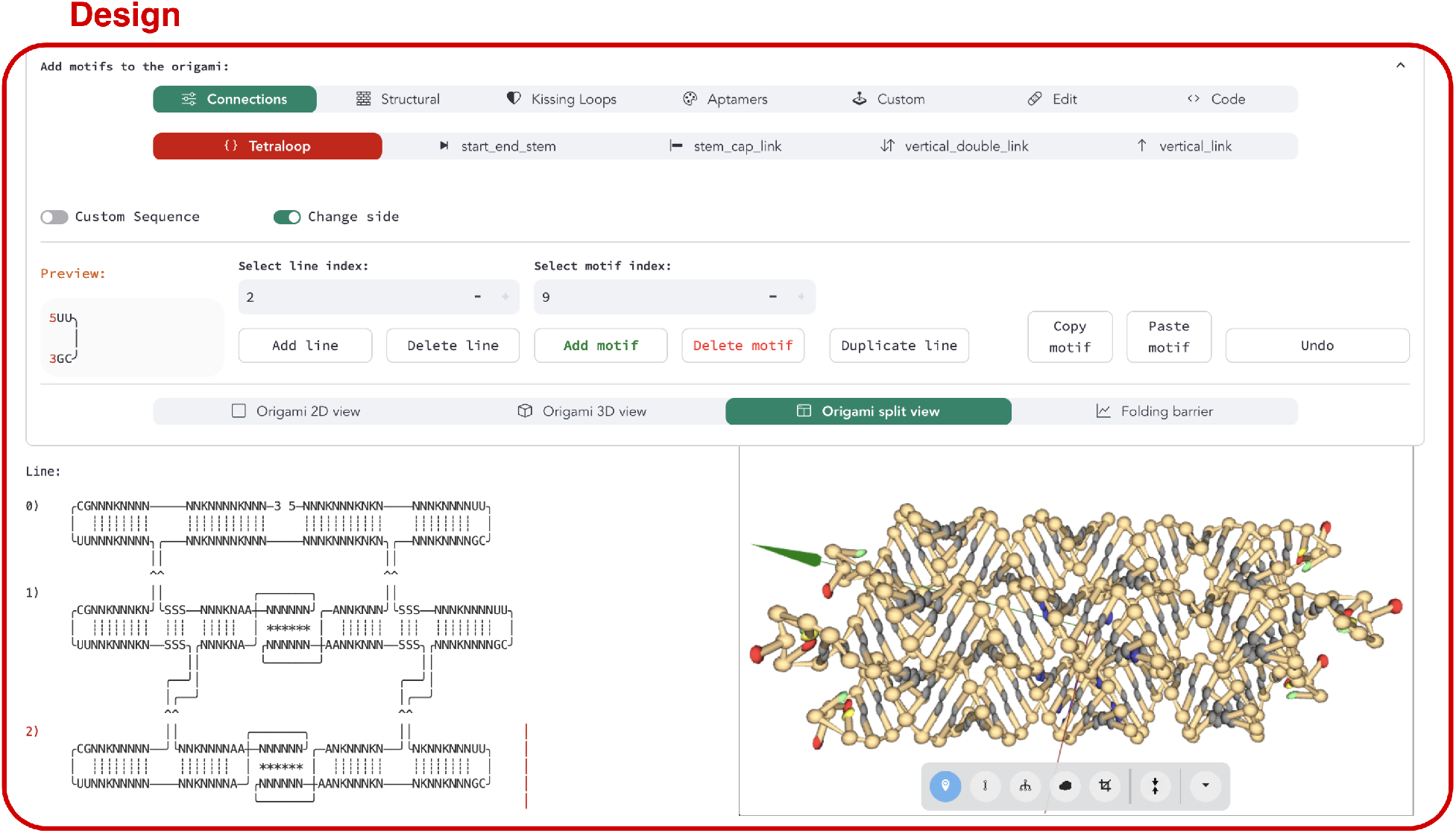
A screenshot of the pyFuRNAce interface (availabe at http://pyfurnace.de). At the top, the motif menu allows for motif selection, creation, and editing via either a graphical or scripting interface. Below, motif properties and a preview are shown, along with index tabs to position the motif within the origami. At the bottom, a menu controls the display settings for the origami; in this case, the origami blueprint is shown alongside the 3D structure in a split view.

Unlike DNA design tools such as CaDNAno [23] or ENSnano [25], which rely on manual strand manipulation, pyFuRNAce adopts a motif-based assembly approach inspired by RNA architectonics and ROAD, extended through object-oriented programming. RNA motifs are collections of strands with assigned structures and 3D representations. With motifs as building blocks, an RNA origami is merely a 2D grid in which the motifs are placed. Users can add, remove, or modify motifs while the software dynamically aligns and connects them, assembling the RNA origami blueprint. At the same time, the origami 3D structure is built by concatenating the motifs’ 3D structures, a feature that enables real-time visualization. Importantly, this also enables the design of structures that consist of multiple RNA strands, which was previously not possible with ROAD [11].

All motifs can be modified, changing the sequences, structures, and 2D representations. Among the numerous motifs available in the RNA nanotechnology field, a relatively small number of structural motifs are sufficient to design an RNA origami nanostructure: tetraloops that cap the end of helices, stems that extend RNA double helices to the desired dimensions, dovetails (stems between crossover junctions) which connect parallel helices at desired angles, and kissing loops, which connect helices within one RNA origami or between multiple origamis (Figure 3a). In pyFuRNAce, these motifs present extra features to allow easy customization of properties such as length, sequence, frequency of wobble pairings or binding energy.

**Figure 3.**
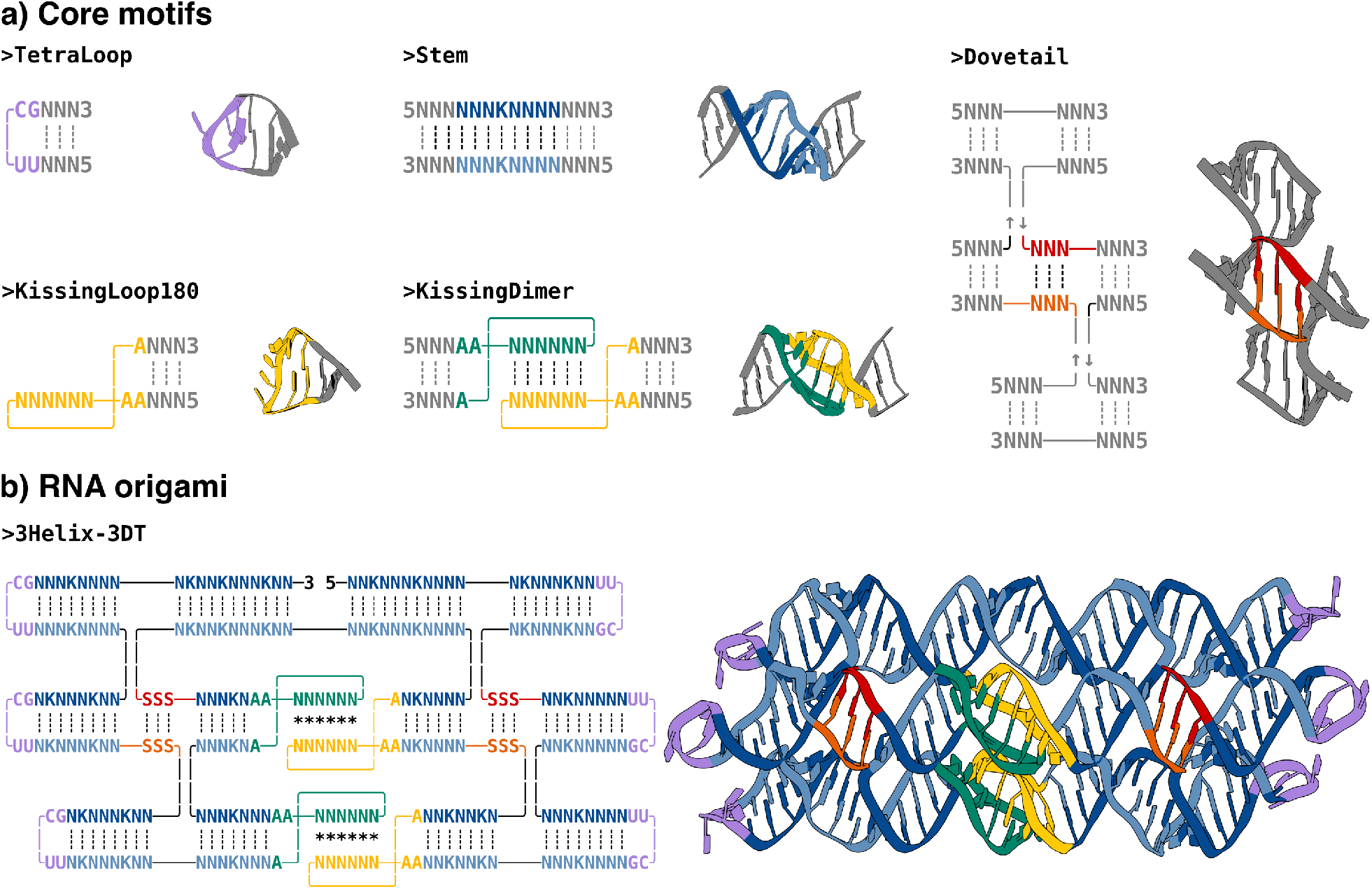
**a)** Most common structural motifs for designing an RNA origami nanostructure. For each motif, the name, blueprint, and 3D structure are shown. Nucleotides that are part of the motif are highlighted in color, while ‘dummy’ motifs are displayed in gray to represent the overall structure, including additional double helices. **b)** An RNA origami blueprint with the corresponding 3D structure. The different motifs in the structure are highlighted using matching colors.

While these motifs are the minimal set to build an RNA origami (Figure 3b), aptamers or ribozymes are necessary to design functional structures. PyFuRNAce includes a growing aptamer library for this purpose (Figure 4a). New aptamers and ribozymes are continuously discovered through experimental techniques such as SELEX (Systematic Evolution of Ligands by Exponential Enrichment) [50] [51] [52] and recently also AI-based tools [53] [54], which promise to tremendously accelerate aptamer discovery in the future. To encourage the expansion of the aptamer library, pyFuRNAce allows users to create custom motifs by defining each strand, their coordinates and 3d structures (Figure 4b). A dedicated custom motif interface is integrated into the design page to streamline this process (in the motif menu, under the option ‘Custom’), with a ‘drawing tool’ tailored towards motif design.

**Figure 4.**
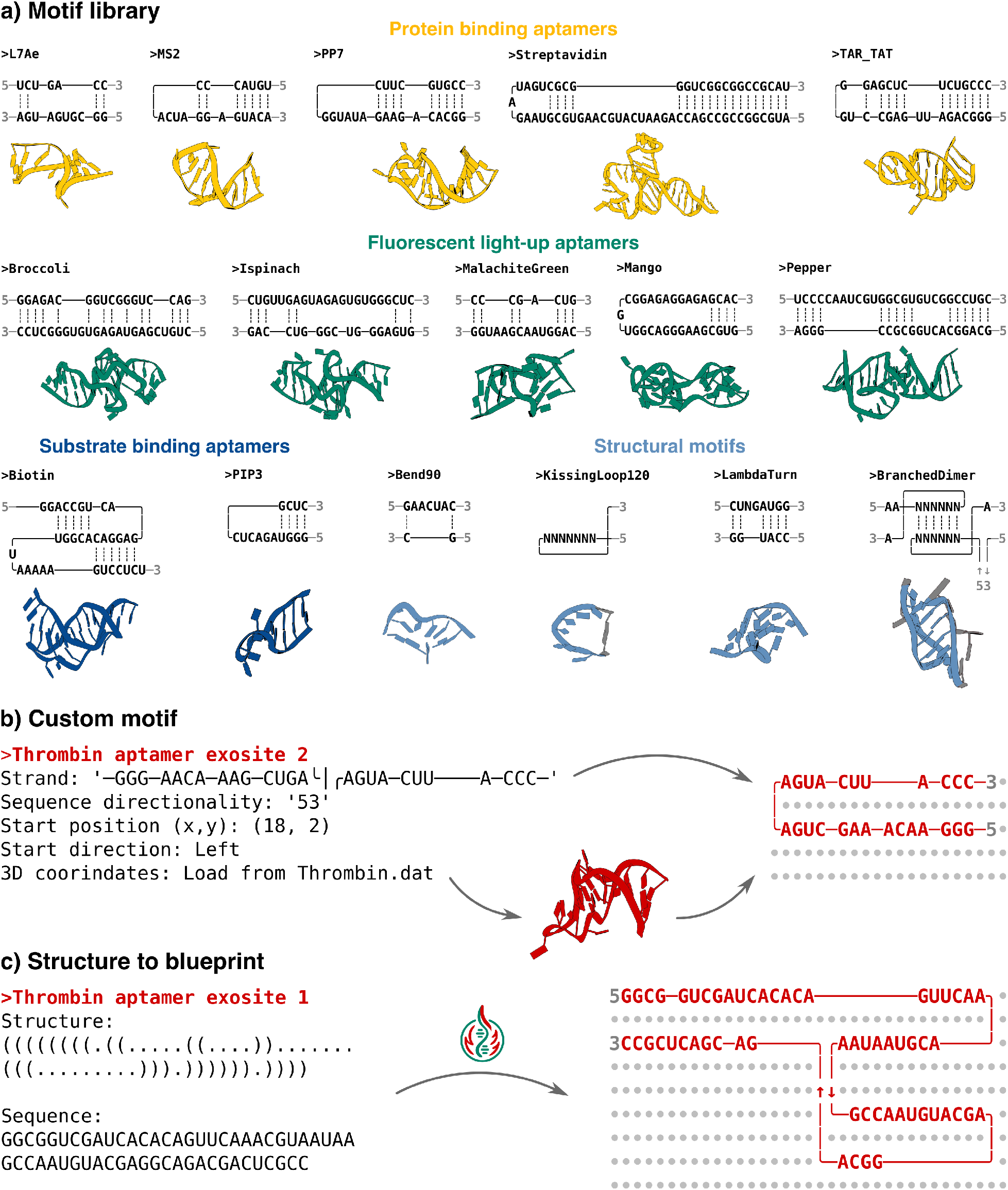
**a)** The current pyFuRNAce motif library (excluding core motifs), with blueprints and 3D structures. The library includes protein-binding aptamers (yellow), fluorescence light-up aptamers (green), substrate-binding aptamers (blue), and structural motifs (light blue). **b)** Custom motif definition: A motif is composed of strands, each defined by strand characters, start position, start direction, and 3D coordinates. The corresponding blueprint is shown on the right. **c)** PyFuRNAce enables the conversion of secondary structure in dot-bracket notation (left) into a corresponding blueprint motif (right), along with an associated sequence.

Recognizing that drawing motifs, either by hand or with drawing tools, can be time-consuming and error-prone, pyFuRNAce includes a Structure Converter tool. This tool converts dot-bracket notation and sequence from FASTA-like files into text blueprints without any need for manual editing. This feature bridges common bioinformatics formats with the pyFuRNAce visual design process, enhancing accessibility and efficiency.

### Generate, Convert, and Prepare

Once an RNA origami structure is complete, the Generate module uses an inverse folding algorithm (Revolvr) to produce an RNA sequence compatible with the target secondary structure. The blueprint is converted into sequence constraints and dot-bracket notation compatible with Revolvr and saved as a target file; then Revolvr is executed in the background. It is important to note that Revolvr requires the installation of Perl and does not yet support kissing loop-specific binding energies. Finally, ViennaRNA folding parameters are provided to evaluate the folding quality, including the energy of the minimum free energy structure, its frequency in the ensemble of possible structures, and ensemble diversity (the average weighted distance between all possible secondary structures).

The Convert module translates RNA sequences into DNA coding sequences and allows the addition of a sequence as a transcriptional promoter at the 5’ end (by default, the T7 promoter is used). It includes analytical tools for determining GC content, dimer prediction, and alignment. The Prepare module supports optional steps such as primer design and oxRNA simulation setup. Primer design is based on melting temperature calculations, with customizable parameters to account for buffer composition and PCR conditions. We recommend cross-validating with melting temperature calculators provided by PCR kit manufacturers. The oxRNA setup includes default simulation files and parameters for evaluating structural stability.

### Programming versatile RNA origami

PyFuRNAce is also a Python package that enables programmable design by scripting. While the GUI lowers entry barriers to RNA origami design, scripting enables advanced customization and control, such as the generation of multi-component structures. For example, a recent RNA origami nanotube design used a 3-helix RNA tile with external 180° kissing loops and a 120° angle between helices[55]. Stem lengths before the kissing loops critically affect tile alignment due to helical twist (due to the periodicity of the helix, one base pair rotates the stem by 32.7°). Previously, alignment was performed manually using 3D visualization tools. With pyFuRNAce, tiles can be programmatically assembled into nanotubes and structural integrity can be evaluated in real time (Figure 5a).

**Figure 5.**
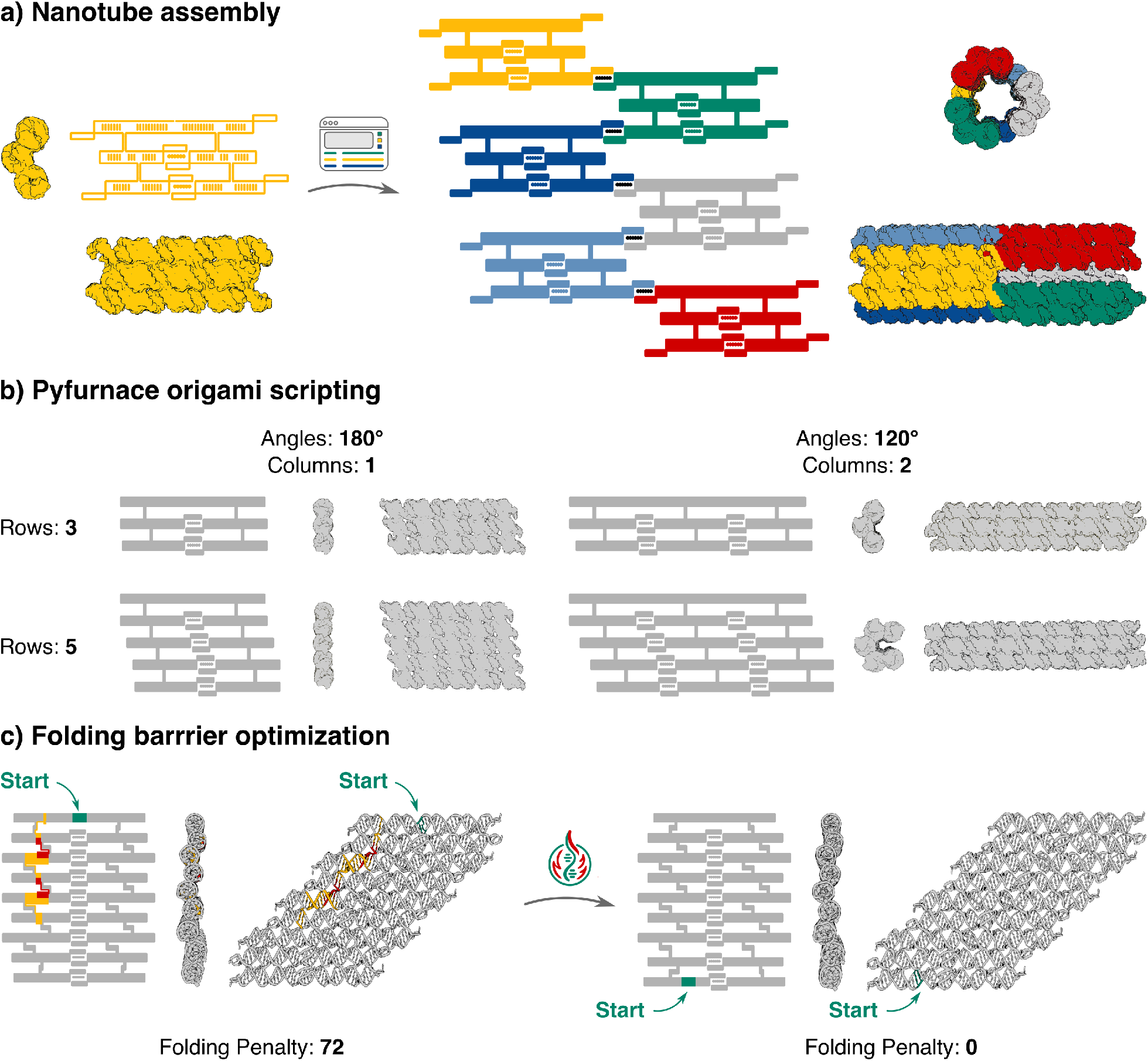
**a)** A blueprint sketch of a single RNA nanotube tile with corresponding 3D structure views on the left, and the assembled nanotube on the right. **b)** Various RNA origami structures that can be generated using the pyFuRNAce Python scripting interface, with customizable parameters such as the number of helices, helix angles, or kissing loop columns. For each structure, the blueprint sketch and two 3D views are shown. **c)** Two equivalent RNA origami structures: one with a moderate co-transcriptional folding barrier (left), where yellow and red colours indicate the severity of the barrier, and an optimized version on the right, with no folding barrier, achieved by rerouting the strand start position.

While multimeric assemblies can reach the micron scale, DNA and RNA origami are often called ‘molecular breadboards’ due to their ability to position single molecules or aptamers with nanoscale precision. For example, different RNA origami structures have been used to precisely position thrombin aptamers as anticoagulants [19]. In particular, RNA origami based on DAE crossovers (double crossover, antiparallel, even spacing)[56, 11] follow predictable design patterns, making them particularly suitable as molecular breadboards. The DAE design rules are: The length of dovetail stems controls the angle between helices; crossover spacing must match the 11-base periodicity of RNA A-form helices; and internal 180° kissing dimers occupy the space of an 8-base stem [47]. Since the pyFuRNAce scripting interface can control all these patterns, we included a “simple origami” function to design DAE RNA origami with ease (Figure 5b). The function requires a few intuitive parameters, such as the number of helices, their angles, or the length of the structure, to generate any DAE design in a few clicks. This capability lets users create customized origami with nanoscale precision while maintaining full control over the individual motifs, and the resulting structures can be downloaded as Python code, ensuring that the design remains flexible and easy to share.

To support co-transcriptional folding, pyFuRNAce integrates the ROAD folding barrier concept: If a pseudoknots pairs before a stem is completely formed, threading of the strand through the duplex is kinetically unfavored and thus is unlikely to form co-transcriptionally. We implemented the barrier checker from ROAD[11], adding an optimization routine that repositions the strand start site to reduce kinetic traps. This produces an equivalent structure with the same overall geometry but a more favourable folding pathway thanks to its optimized sequence (Figure 5c).

These features aim to accelerate RNA origami development, improve design reliability, and make advanced design strategies more accessible to a broader community.

## Discussion

RNA origami is a powerful method for designing functional and programmable RNA nanostructures with wide-ranging applications in nanomedicine, synthetic biology, and biosensing. Due to the high stability of RNA structures and the possiblility for multiplexing of e.g. aptamers, we see tremendous potential in the use of RNA origami for therapeutic applications. However, the lack of integrated and user-friendly design tools has hindered the broader adoption of RNA origami.

PyFuRNAce addresses this gap with several key contributions to the landscape of RNA origami design software:

1. *Integrated graphical interface:* pyFuRNAce offers an intuitive, fully integrated graphical user interface that lowers the entry barrier for new users while providing advanced features for experienced designers. The interface enables real-time updates and seamless transitions across design stages with a few clicks, simplifying a process that previously required multiple disjoint tools.
2. *Custom motifs support:* The rapid discovery of new functional RNA motifs, e.g. through SELEX and enhanced by AI, demands a flexible design environment. PyFuRNAce allows users to define and integrate new custom motifs, including aptamers and ribozymes, integrating the latest structural innovations into RNA origami designs.
3. *Programmability:* While interactive design is essential, some tasks—such as defining scaffold layouts, testing structural variants, or optimizing folding pathways—require extensive user inputs or manual modification. PyFuRNAce’s Python interface allows users to build complex structures by scripting, accelerating the design process and enabling high-throughput applications.

Despite these advantages, pyFuRNAce has some current limitations. Its blueprint representation is optimized for adjacent motifs, and non-linear connections (e.g., between non-adjacent helices) are not yet straightforward to design. Additionally, the sequence generation engine (Revolvr) lacks support for multi-strand optimization and fine-grained control over individual kissing-loop energies. To address these issues, we are developing a new inverse folding algorithm tailored to RNA origami, to expand pyFuRNAce’s sequence design capabilities to match its structural design flexibility. We still believe that pyFuRNAce in its current form is a useful tool which will help grow a larger community around RNA origami. This is why we chose to already share it in its current form.

In summary, pyFuRNAce provides a comprehensive and accessible platform for RNA origami design, bridging the gap between abstract structural modelling and practical sequence implementation. Integrating design, visualization and sequence optimization into a single environment, pyFuRNAce empowers researchers from diverse fields to easily design RNA nanostructures with speed, accuracy, and creativity.

## Software availability

The pyFuRNAce source code, documentation, and examples are available on Github at the repository: https://github.com/Biophysical-Engineering-Group/pyFuRNAce. The package can be installed via the Python Package Index (PyPI) using the standard Python package installer (pip). The web application is hosted on Streamlit Cloud and is freely accessible at: http://pyfurnace.de.

## Acknowledgements

This work was supported by the ERC Starting Grant “ENSYNC” (No. 101076997) and Deutsche Forschungsgemeinschaft (DFG, German Research Foundation) under CRC 392 and CRC 1638. K.G. acknowledges funding from the Human Science Frontier Programme (HFSP). The authors thank the Max Planck Society for access to computational resources and the Alfried Krupp von Bohlen und Halbach Foundation. We thank C. Geary for his valuable input on the manuscript and insightful feedback on the user interface design.

